# The compartmental polarization of interphase human chromosomes within the nucleus: A mechanism for transcriptional regulation of large chromosomes

**DOI:** 10.64898/2026.08.21.746281

**Authors:** Matheus F. Mello, Dimanthi N. Perera, Lauren E. DiPaola, Antonio B. Oliveira, Vinícius G. Contessoto, José N. Onuchic, Ryan R. Cheng

## Abstract

The structure and dynamics of individual chromosomes are significantly modulated by their local environment within the nucleus. It has long been established that heterochromatin (B compartment loci) associates with the lamina meshwork that lines the inner nuclear envelope. However, the extent to which these interactions perturb the chromosomal organization within a territory remains unclear. Using computer simulations and published imaging data, we further characterize the interaction between chromatin and the lamina and find striking consequences of lamina-association on the chromosomal structural ensemble. We find that lamina-associating chromosomes have their compartmentalization polarized in a direction perpendicular to the nuclear surface. Further, lamina-associated chromosomes exhibit shorter average distances between euchromatin loci (A compartment) than chromosomes without lamina contact. Energy landscape analysis of our simulations reveal that the sequestration of heterochromatin to the lamina allows for euchromatin to form more spatial contacts with other segments of euchromatin. The compaction of euchromatin due to lamina-association also diminishes the mobility of these regions. These findings suggest a mechanism for bringing together euchromatic segments that are separated by large genomic distances within a chromosome, potentially to share transcriptional machinery and enhance transcription for large chromosomes, which are often found at the nuclear periphery.

**Significance statement:** The structures of human chromosomes are highly dynamic and influenced by their local environment within the nucleus. A key environmental constraint is given by the nuclear membrane itself. The inner nuclear membrane is lined with protein filaments called the nuclear lamina, which directly interact with segments of inactive heterochromatin. We use published experimental imaging data combined with physics-based polymer models to investigate how lamina-association influences the three-dimensional organization of chromosomes. We find that the sequestration of inactive heterochromatin to the lamina promotes compaction of active euchromatin away from the lamina, similarly to allostery in proteins. Our results suggest that association with the nuclear periphery may enhance transcription by bringing actively transcribed regions into closer proximity, potentially facilitating the sharing of transcriptional machinery and splicing factors.

## I. INTRODUCTION

The spatial organization of the genome plays a central role in its function, maintenance, and regulation. In eukaryotic cells, individual chromosomes typically form distinct territories within the nucleus, as seen in the famous Cremer and Cremer microscopy image [1]. These chromosomes are further organized through micro-phase separation of active (euchromatin) and inactive (heterochromatin) chromatin [2]. The two compartments resulting from the segregation of euchromatin and heterochromatin are referred to as compartments A and B, respectively. In addition, chromosomes are shaped by interactions between heterochromatin and the nuclear lamina [3]. However, a comprehensive picture of genome organization and its relationship to biological function remains elusive.

Recent developments in the theoretical understanding of chromosomal organization have shown that chromatin structure can only be thought of from an ensemble perspective [4, 5]. This has also been supported by high-resolution DNA tracing experiments (microscopy) [6–9] and single cell Hi-C [10]. At the level of individual territories, chromosomes are organized by chromatin compartmentalization, as well as lengthwise compaction facilitated by the activity of condensin and cohesin [2, 11–13]. These physical considerations were used to construct a theoretical framework for genome organization called the Minimal Chromatin Model (MiChroM) [5, 14]), which is a coarse-grained polymer model of chromosomes. The structural predictions of MiChroM have been shown to be quantitatively consistent with Hi-C [5, 14, 15] and fluorescence imaging [15], capturing not only structural organization but also viscoelastic dynamics [16] and response to mechanical tension [17].

Certain heterochromatic segments have been found to associate with the nuclear lamina [3, 18, 19]. This further acts as a significant constraint on the spatial organization of the chromosomes [20], and down-regulation of the lamina proteins results in a “nuclear inversion”, where the dense heterochromatin is organized in the interior of the nucleus [21]. Numerous studies have also shown that the proximity of the chromosome to the nuclear envelope appears to be positively correlated with chromosome size [22] and negatively correlated with gene density [23]. This size-dependent proximity has also been shown to directly influence the spatial neighborhood of individual chromosomes − i.e., the chromosomes or nuclear bodies that are adjacent to an individual chromosome. Understanding the effects of proximity to the nuclear periphery on chromosome organization and dynamics is the core focus of this study.

Technological advances (i.e., DNA tracing [6–9]) have further allowed chromatin structures to be imaged from individual cell nuclei using a microscope. Although early tracing experiments have focused on short segments of chromatin, recent advances have led to imaging of the entire human genome [8]. These experiments have generated thousands of individual snapshots of chromatin structures within the nuclei of a population of cells.

In this study, we combine available experimental data of genome-wide traced structures [8] with a theoretical model of the nucleus to decode the role of the lamina on a chromosome structure and dynamics. This model consists of a chromosome described by MiChroM confined within a spherical nucleus, subject to interactions with the lamina and nucleolus. We find that the nuclear lamina has a significant perturbative effect on the structural ensembles of the chromosomes. A key finding is that the nuclear lamina serves to polarize the compartmental microphase separation of chromosomes, resulting in the compaction of the A compartment. Although the sequestration of B compartment loci to the nuclear lamina is well established [3], we report that a major impact of this sequestration is that A compartment loci are able to form more contacts with other A loci (i.e., the average A-A distances are shorter and more compact). This increased compaction of the A compartments further decreases the mobility of these segments. Our results also suggest that the perturbative role of the nuclear lamina is more pronounced for larger chromosomes than for smaller ones. The work highlights how even subtle changes to the spatial positioning of a chromosome can result in structural changes across an entire territory, offering a potential mechanism to enhance transcription for large chromosomes by bringing together active segments or modulate the function of chromosomes through small perturbations in nuclear positioning.

## II. RESULTS

### A. Spatial polarization of compartmentalization due to chromosome contacting lamina

The spatial positioning of interphase chromosomes within a nucleus is not random; larger chromosomes are more frequently observed to be in contact with the nuclear periphery [22, 23]. This is further observed in imaged fibroblast cells [8], where larger chromosomes, often rich in heterochromatin content, have a high probability of interacting with the lamina (Fig. S1A). It has further been established that B compartment chromatin associates with the nuclear lamina in the periphery of the nucleus [3]. However, what has not been thoroughly discussed is the perturbative role of lamina-association on the structural ensembles of the chromosome territories.

For chromosomes contacting the lamina, the sequestration of B loci to the lamina acts as a significant constraint on the structural ensemble. This constraint would be expected to cause (on average) an enrichment of A loci in the perpendicular direction moving away from the lamina.

To quantify this effect, we propose a compartment polarization defined as

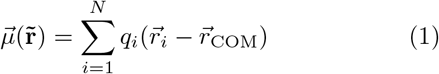

where *i* represents each annotated locus in the chromo-some (i.e., not assigned as NA), and

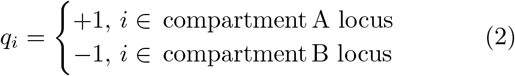

We computed the compartment polarization for each chromosome in the genome-scale imaging dataset [8] (Fig. 1A,B). For a given chromosome, the compartment polarization captures an axis along which compartments A and B spatially segregate. For each cell, the nuclear envelope can be estimated by calculating a convex hull around the imaged chromatin loci [8]. We consider a locus to interact with the lamina if it belongs to the hull boundary. By extension, a chromosome is considered to interact with the lamina (∈ lamina) when 10% or more of its loci in-teract with the lamina, and not to interact (*∉* lamina) when none of its loci is found in the hull boundary. In addition, we use the convex hull to estimate an effective normal vector of the nuclear surface based on the local boundary region contacted by or nearest to each chromosome (see Appendix A). Fig. 1A illustrates this compartment polarization 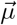 relative to the normal vector to the nuclear surface 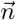. When the chromosomes interact with the lamina, we observe a shift in the distribution of the angle formed by the compartment dipole and the direction of the lamina towards alignment of these two vectors (Fig. 1C). Given the presence of many other influences in the nucleus, such as other chromosomes and nuclear bodies [24, 25], the difference between the distribution of the alignment angle between the dipole and the lamina shows a tendency for the lamina to polarize the configuration of the compartments inside the territory.

**FIG. 1.**
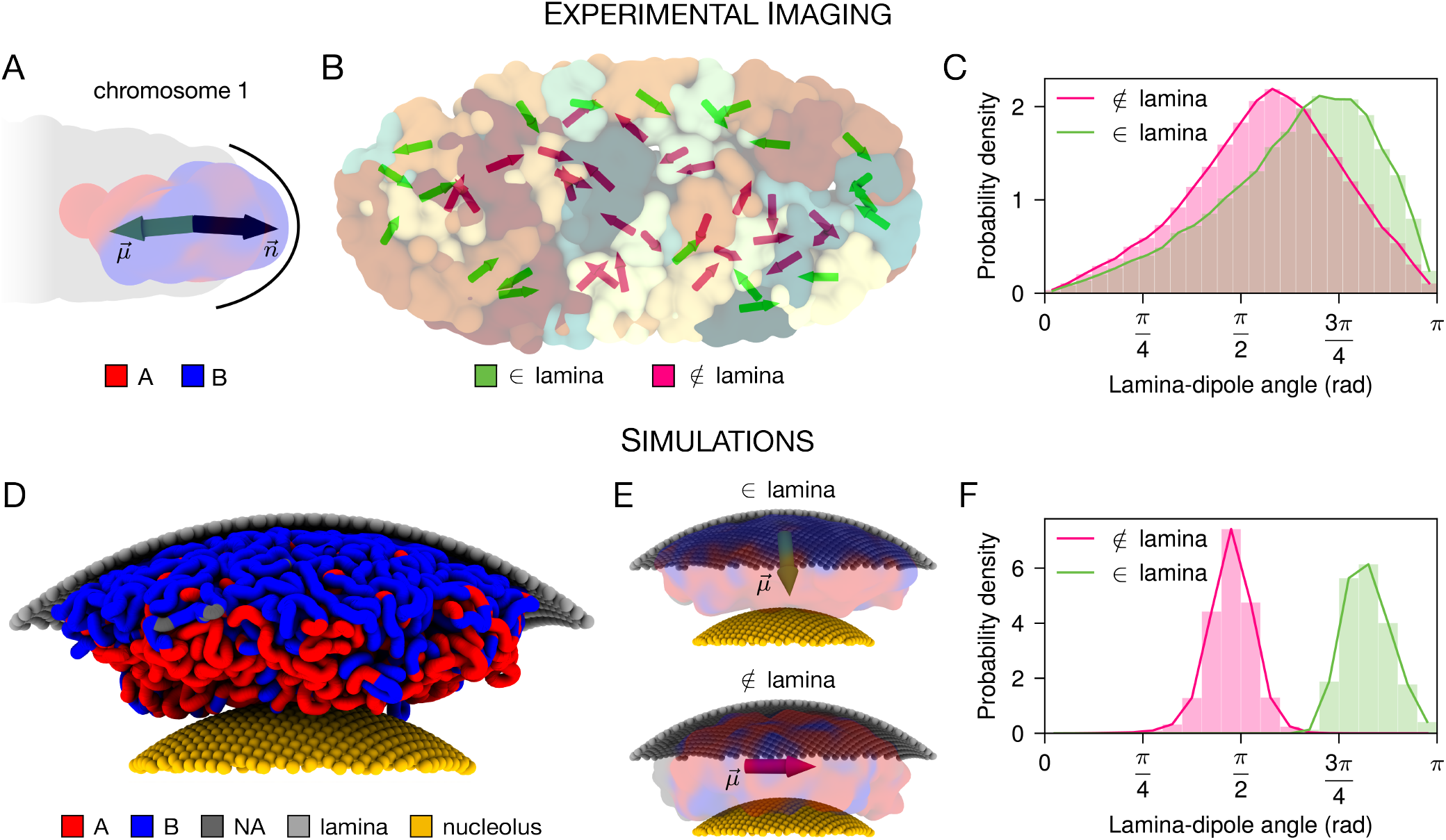
Compartmental polarization and its alignment with the nuclear lamina. (**A**) Representative structure of chromosome 1 in the nucleus as imaged by Ref. [8]. The chromosome is colored by compartments, as annotated by MEGABASE for IMR-90. The compartment polarization 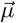 is represented in green, together the normal vector 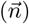 to the nuclear periphery, as estimated by the whole genome convex hull. (**B**) Representative structure of the nucleus as imaged by Ref. [8]. For each chromosome territory, the direction of the compartment polarization is shown. When the chromosome is in significant contact with the lamina, i.e., 10% or more loci defines the nuclear convex hull, its dipole is colored green, and magenta otherwise. (**C**) Probability density distribution of the angle between the vectors of the compartmental dipole and the normal to the lamina for chromosomes interacting (green) and not interacting (magenta) with the lamina. When a chromosome does not touch the lamina, i.e., no imaged loci for that chromosome defines the nuclear convex hull, we use the normal on the closest point on the periphery. (**D**) Representative structure of the molecular dynamics simulation of chromosome 1 of human cell line IMR-90, with the Minimal Chromatin Model (MiChroM) [5]. The chromosome is colored by compartments, as annotated by MEGABASE for IMR-90. The chromosome is positioned between the nuclear periphery and the nucleolus. (**E**) Representative structure of simulated chromosome 1 in the presence (top) and absence (bottom) of chromatin interactions with the lamina. The direction of the compartmental dipole is shown together with the surface of the chromosome in each case. (**F**) Probability density distribution of the angle between the vectors of the compartmental dipole and the normal to the lamina for the simulated chromosome 1 in the presence (green) and absence (magenta) of chromatin interactions with the lamina.

To probe the extent of this compartment polarization in the absence of other interacting chromosomes, we employed the Minimal Chromatin Model (MiChroM) [5] together with MEGABASE [14] to simulate chromosome 1 of human cell line IMR-90 at a resolution of 50 kb, with (∈ lamina) and without (∉ lamina) attractive interactions between the chromatin and the lamina. The chromosome is located between the nuclear periphery and the nucleolus (Fig. 1D). The strength of interaction with the lamina and the nucleolus was determined based on the differences between the contact probability of B versus A loci in the imaged ensemble (see Appendix B for details on simulation methods). Similarly to the results observed for the imaging structures, the compartmental polarization of the simulated chromosome aligned with the lamina direction only when lamina interactions were present (Fig. 1E,F). Both experimental and simulated distributions exhibit similar lamina-dipole angles, illustrating the role of the nuclear lamina in polarizing chromatin compartmentalization in a territory. It should be noted that although the experimental and simulated distributions present peaks with similar dipole angles (Fig. 1C,F), the experimental distributions are notably broader compared to simulation, likely due to the crowded nuclear environment that includes other chromosomes and nuclear speckles, which are not considered in the simulation model.

### B. Sequestration of B loci to lamina results in compaction of A compartment within a territory

Previous studies have reported that although the A/B compartment assignment remains mostly the same, some changes in the cis contact frequencies (i.e., for loci in the same chromosome) are observed when chromosomal interactions with the lamina are disturbed [26, 27]. By annotating which chromosomes contact the nuclear periphery (or do not) in each cell in the imaged structural ensemble [8], we can analyze how interactions with the lamina affect the distance between pairs of chromatin loci without requiring additional experimental data where lamin proteins are depleted. Therefore, we selected cells in which the chromosome did not contact the nuclear envelope (*∉* lamina) and cells in which 10% or more of the imaged loci were part of the periphery (∈ lamina). Although limited by the number of imaged loci per chromosome, we observed that chromosomes exhibit a significant shift in their pairwise distance distributions towards smaller distances upon interaction with the lamina (Fig. 2A and Fig. S2). To better understand this shift and its relationship to genomic distance, we further examined the bi-variate distribution of the pairwise distance and genomic distance between pairs of imaged loci within an individual chromosome (Fig 2A). We observe an enrichment of the probability density for the ∈ lamina condition relative to the *∉* lamina condition for shorter pairwise distances ranging from *∼* 2-3 *µ*m across genomic distances. This behavior was observed to be stronger for larger chromosomes (Fig. S2) and more pronounced for A-A pairs than for B-B or A-B pairs (Fig. S3).

**FIG. 2.**
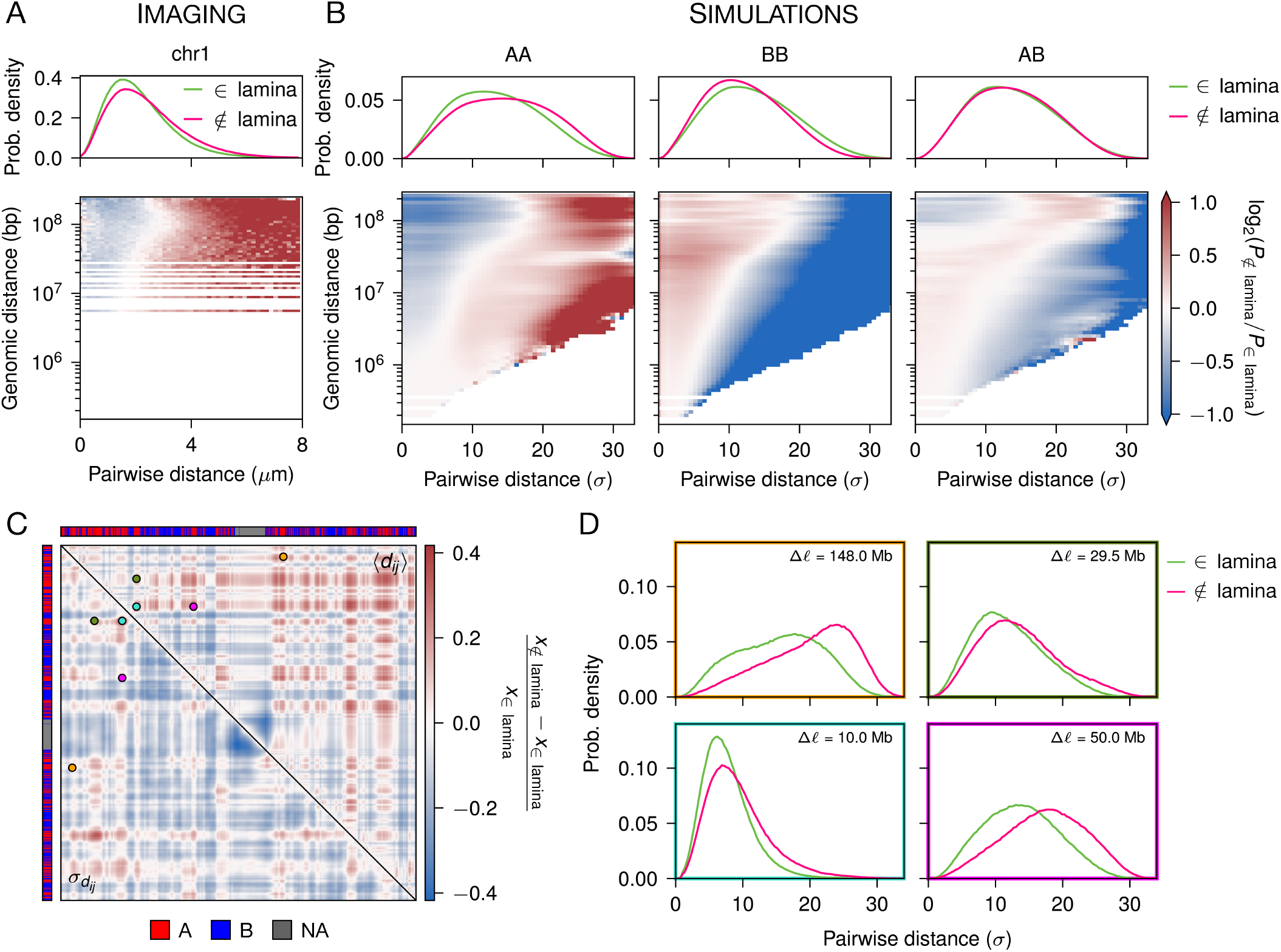
Lamina-mediated compaction of euchromatic regions in experimentally imaged and simulated structural ensembles. (**A**) For the homologs of chromosome 1 in the whole-genome ensemble imaged by Ref. [8], pairwise distance distributions when the chromosome interacts with the lamina or not (top) and log ratio between the bi-variate probability density distributions P_∉ lamina_(d_*ij*_, r_*ij*_)/P_*∈* lamina_(d_*ij*_, r_*ij*_) (bottom), where d_*ij*_ and r_*ij*_ respectively represent the genomic and spatial distances between loci i and j. (**B**) Same as (A) for the simulated chromosome 1, separated by type of pair: AA, BB, and AB. (**C**) Relative difference of the average pairwise distance matrix (top diagonal) and of the standard deviation of the distance matrix (bottom diagonal) comparing the simulated ensembles without and with lamina interactions. Positive values indicate higher compaction when the chromosome interacts with the lamina, while negative values indicate higher compaction in the case without those interactions. (**D**) Pairwise distance distributions for selected AA pairs highlighted in (C). When the chromosome interacts with the lamina, there is an enrichment of smaller distances for AA pairs.

The computational model of chr1 allows us to better understand this perturbative role of the lamina on the chromosomal structural ensembles. We find a more pronounced difference between the statistical behavior of A-A and B-B chromatin pairs in the simulated ensemble for ∈ lamina compared to the *∉*lamina condition. In particular, A-A pairwise distances are shorter for ∈ lamina simulations, indicating a compaction of the A compartmentalization upon the sequestration of B loci to the lamina (Fig. 2B). This shortening of A-A distances broadly affects pairs of loci separated by genomic distances from 10^6^ bp to the length of the chromosome (Fig. 2B), and results in a significant relative shift in the bi-variate probability density towards shorter distances for A-A loci pairs separated by *∼* 10^7^-10^8^ bp for ∈ lamina relative to ∉ lamina. Although the imaging data has significantly lower resolution than the simulation, there is quantitative agreement between the simulation and the experiment, when considering the conversion from reduced to physical units (1 *σ ≈* 0.14 *µ*m [14, 28]). For example, the transition from a negative to positive log_2_ ratio around 10^8^ bp occurs between *∼* 2.5-3.4 *µ*m for the experimental data, which is in a similar range obtained for the transition of A-A pairs in the simulations (*∼*17-19.2 *σ ≈* 2.4-2.7 *µ*m).

On the other hand, the opposite trend is observed for compartment B (Fig. 2B). B-B pairwise distances were on average longer when the chromosome interacts with the lamina. This decompaction of B compartments, together with the compaction of the A compartments, can also be observed in Fig. 2C, which shows a representation of the relative deviations between the mean distance matrix and its standard deviation of the *∉* lamina vs. ∈ lamina simulation ensembles. A-A pairs are on average closer in the presence of lamina interactions, and their distance distributions have a smaller standard deviation (Fig. 2C). This can be seen more explicitly in Fig. 2D, which compares the pairwise distance distributions of A-A pairs separated by different genomic distances for the two simulated conditions.

Together, these results indicate that interactions with the lamina lead to a higher compaction of the A compartments, offering a mechanism for transcriptional regulation − e.g., proximal segments are able to better share transcription machinery.

### C. Energetic stabilization of A compartment and destablization of B compartment by the lamina

To understand the underlying cause of these shifts in the distance distributions, we investigated the energet-ics of the simulated trajectories. When comparing the ensembles of structures with and without interactions with the nuclear lamina, we observe that the total contact energy is lower by approximately 0.12 *k*_B_*T* per bead when the chromosome is in contact with the lamina (Fig. 3A). For comparison, the maximum energetic contribution of a chromatin–chromatin contact ranges from 0.23 to 0.33 *k*_B_*T*, depending on the compartment annotations of the interacting beads, while the actual contribution depends on their distance (see Appendix B). Most of this stabilization comes from the added interaction with the lamina (Fig. 3B). We also observe that the total contact energy of B-B loci is smaller when lamina inter-actions are present, suggesting that some B-B contacts are destabilized in order to establish these interactions with the lamina (Fig. 3C). On the other hand, contacts within compartments A are further stabilized when lamina interactions are present (Fig. 3D).

**FIG. 3.**
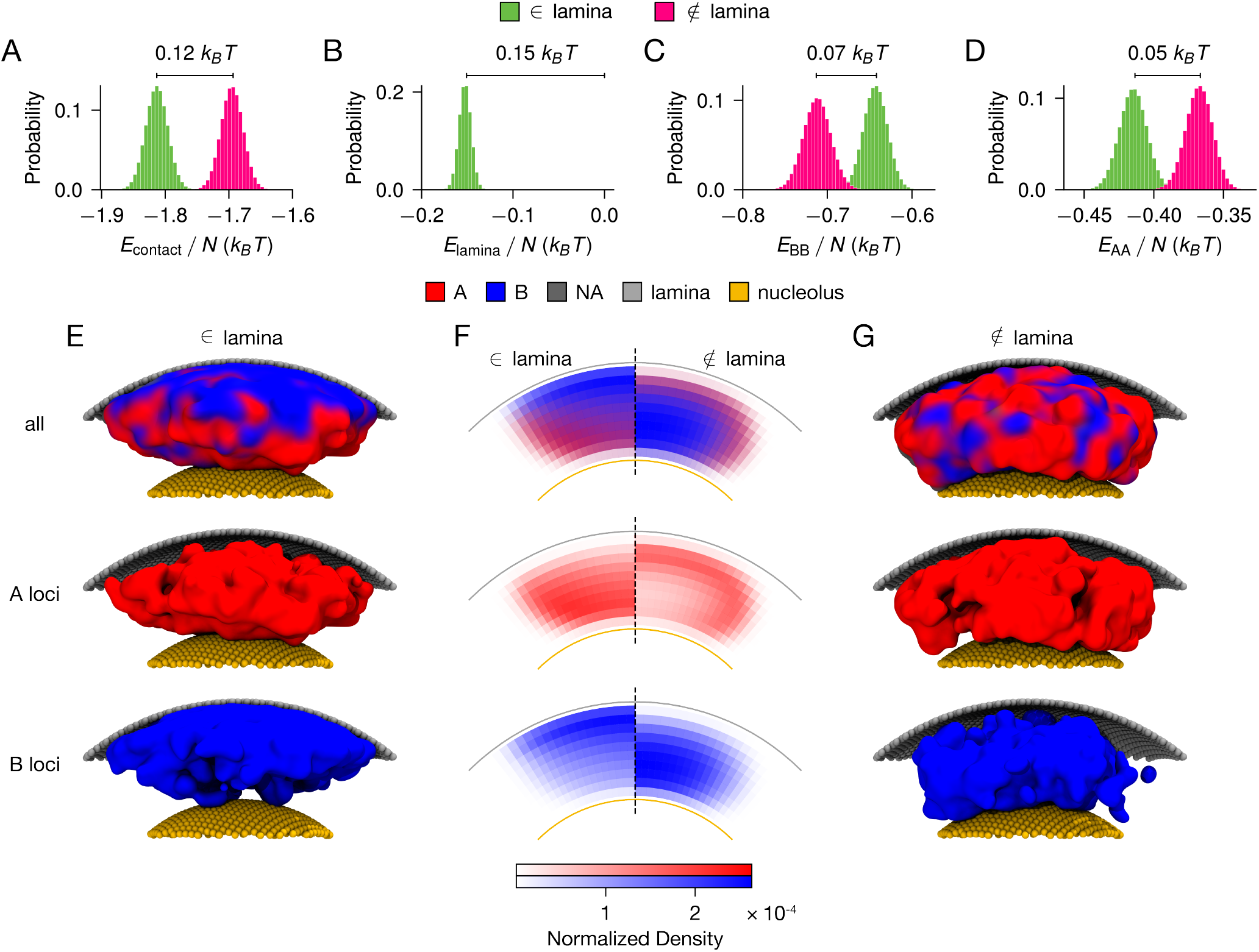
Interactions with the lamina further stabilize compartment A. (**A**-**D**) Probability density distributions for contact energies, E, normalized by the number of beads of the chromosome, N, across the trajectories with (∈ lamina) and without (∉ lamina) chromatin interactions with the nuclear periphery. (**A**) Total contact energy, which includes chromatin-chromatin interactions, as well as chromatin interactions with the lamina and the nucleolus. (**B**) B-Lamina contact energy. (**C**) B-B contact energy. (**D**) A-A contact energy. (**E**) Representative structure of the ensemble with chromatin-lamina interactions. The whole territory is shown (top), together with representations including only the A (middle) or B (bottom) loci. (**F**) Normalized density profile, ρ(r, θ), computed over the structural ensemble for each condition. Each (r, θ) bin represents the relative volume density after integration over the azimuthal angle. For each condition, these maps provide a statistical description of the organization of each compartment within the territory. (**G**) Representative structure of the ensemble without chromatin-lamina interactions. The whole territory is shown (top), together with representations including only the A (middle) or B (bottom) loci.

In the absence of external interactions, the chromosome [5] and the nucleus [21, 29] are organized with a core of inactive chromatin, while active loci occupy the periphery of the territory or nucleus, respectively. As B-B interactions are energetically favored compared to A-A [5, 14, 29], the formation of this central cluster of B-type loci minimizes the free energy of the system, as it maximizes B-B contacts [30]. Upon additional interactions, such as contacts with other chromosomes or nuclear landmarks, this theoretical organization of the territory is perturbed [31]. As a consequence, the composition of the compartments on the surface of the chromosome changes.

In our case, when lamina interactions are considered, we notice an increase in B-type loci that occupy the lamina-exposed surface of the chromosome (Fig. 3E-F and Fig. S4A). However, the B-compartment contact network remains largely unchanged (Fig. S4B), with the majority of inactive chromatin participating in a core contact group. As a consequence of the sequestration of the B loci to the nuclear periphery, A-type loci populate the outer layers of the internal “hemisphere” of the chromosome territory (Fig. 3E,F). This joint effect of geometrical and energetic constraints, where the B compartment is brought to the nucleus periphery while maintaining most of its energetically-favorable contacts (Fig. 3C), reduces the conformational space of active chromatin, increasing the likelihood of A–A contacts.

### D. Affect of lamina-association on dynamics of chromosome

We further examined the computer simulations to understand the role of the nuclear lamina on the conformational dynamics of chromatin. In Fig. 4A, the calculated mean squared displacement (MSD) shows that the mobility of A compartment loci is enhanced for the∉lamina condition. On the other hand, the overall mobility of B loci appears unaffected by the interaction of the chromosome with the lamina. Note that this measurement represents an average behavior that involves loci interacting with the lamina and loci located more internally in the chromosome territory.

**FIG. 4.**
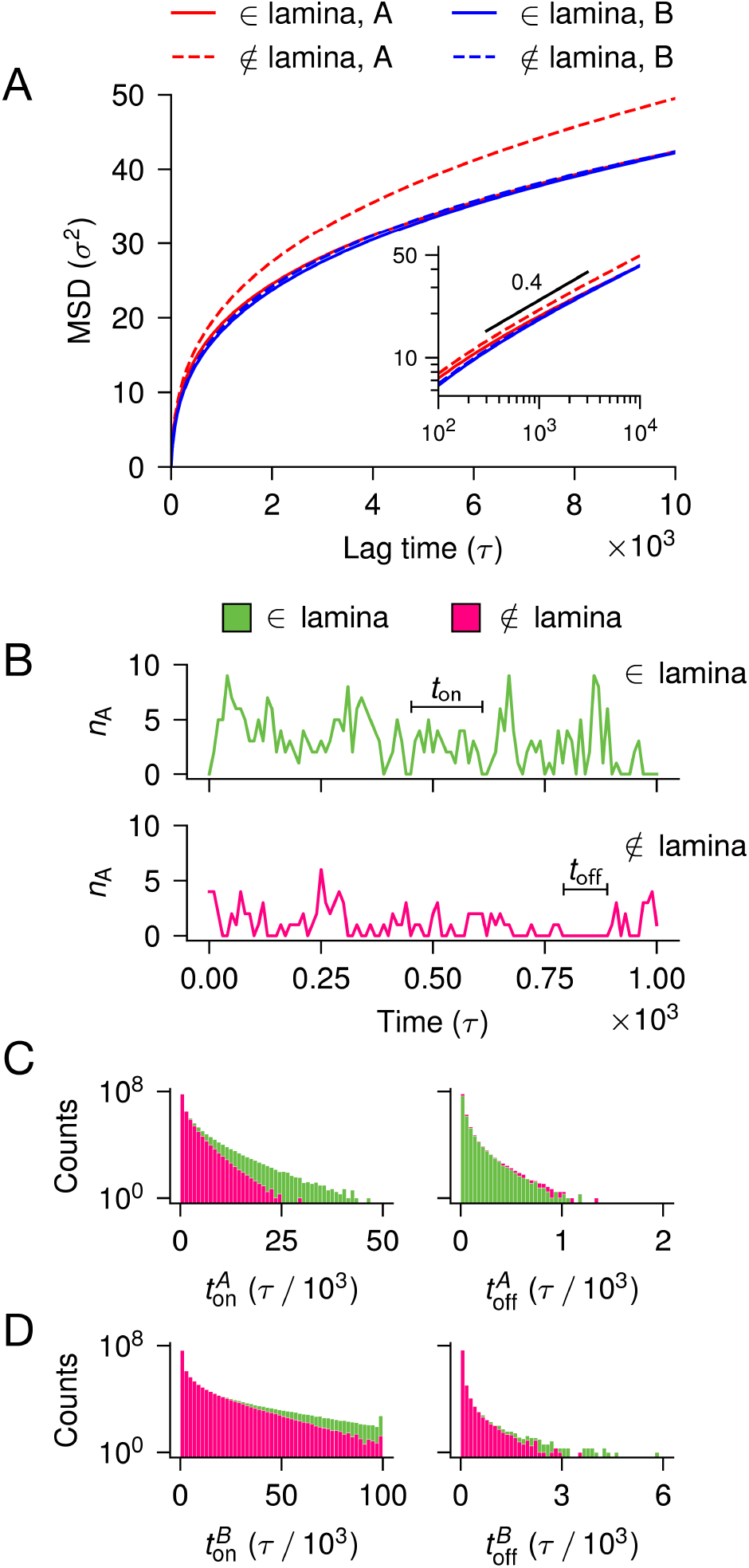
Lamina effect on the dynamics of the model. (**A**)Mean squared displacement (MSD) of A and B loci for the structural ensembles with and without chromatin interactions with the nuclear periphery. The inset shows the same data in log scale, together with the slope of 0.4. (**B**) Time traces of the number of contacts with non-neighboring A loci for a particular A locus, in the trajectories with and without lamina interactions. From the time traces, we measure the duration a locus stay in contact with any other loci of the same compartment (t_on_) and the duration it stays without any contact with any other loci of the same compartment (t_off_). (**C**) Distributions of t_on_ and t_off_ for A loci in the trajectories with and without lamina interactions. (**D**) Same as (C), but for B loci.it is likely to be consistently bound [32].

In addition, we quantified the temporal behavior of compartmentalization by characterizing *t*_on_, the duration of time a locus stays in contact with any other loci of the same compartment, and *t*_off_, the duration of time a loci stays without any contact with any other loci of the same compartment. Fig. 4B shows a representative time trace for each condition, with examples of *t*_on_ and *t*_off_. We find that for A compartment loci, the ∈ lamina condition results in longer *t*_on_ times than for the *∉* lamina condition, while *t*_off_ distributions are not affected (Fig. 4C, Table S3).

For the B compartment loci, the interactions with the lamina also subtly increase the probability of long-forming contacts (Fig. 4D), although the average *t*_on_ remains the same in both conditions (Table S3). Interestingly, this happens even with a smaller B-B contact energy for the ∈ lamina condition (Fig. 3). As the lamina provides a geometric anchoring for the B loci (Fig. 3E-G), B loci are more likely to be within at least one non-neighboring contacting locus, even if they do fewer contacts overall (Fig. S4).

Our simulation results are consistent with previous experimental observations [27, 32]. It has been shown that when lamin B1 is knocked out in human cells, regions of the chromosome that are consistently found in the nucleoplasm have enhanced dynamics [27], suggesting that indeed chromatin-lamina interactions affects the behavior of the whole territory, instead of being a more localized interaction in the vicinity of the territory/lamina region. Another interesting experimental result shows that binding to the lamina is usually cooperative for lamina-associated domains (LADs) on the same chromosome, suggesting that indeed if a chromosome has more LADs,

## III. DISCUSSION

In this work, we have coupled experimental and simulated data to investigate how interactions with the nuclear lamina shape the organization of chromosome territories. In particular, we analyzed the organization of compartments within the chromosome. We observed in both imaged and simulated ensembles of structures that the interaction with the lamina creates an axis in the direction normal to the surface in which compartments A and B segregate.

Interestingly, proximity to the nuclear periphery not only changes the relative positioning of the compartments within the chromosome territory but also shapes the chromatin organization within the compartments. For chromosomes with lamina interactions, compartments A and B are, respectively, more and less compact compared to chromosomes that lack these interactions. The compaction of the A compartment chromatin resulting from the sequestration of B chromatin to the lamina offers a potential mechanism for transcriptional regulation, particularly for large chromosomes that are often observed proximal to the lamina.

Large chromosomes, such as chromosome 1, have been noted to be heterochromatin rich and are frequently observed to be associated with the lamina [33]. The compaction of a chromosome due to lamina-association allows A compartment loci not only to form more contacts with other A loci and to be on-average closer together. This enhanced spatial proximity of A compartment loci to one another can potentially allow active genes located on those loci to share, for example, transcriptional machinery such as RNA polymerase and splicing factors.

Our results potentially offer a mechanism for diseases that may disrupt the compartmental polarization or the compaction of lamina-associated chromosomes, such as certain large translocations associated with cancer [34] or epigenetic misregulation associated with disease states [35]. Further, the results discussed here may also play an important role in the mechanosensory role of the nucleus, as mechanical perturbations subtly shift the organization of the genome [36, 37].

## ACKNOWLEDGMENTS

We thank Dr. Sumitabha Brahmachari for many useful conversations during the development of this work and for all his comments and suggestions. RRC would also like to thank Jason Parsley and Eric J. Rawdon for helpful discussions. RRC acknowledges support from NSF Award PHY-2540931 and acknowledges the University of Kentucky Center for Computational Sciences and Information Technology Services Research Computing for their support. Work at the Center for Theoretical

Biological Physics was supported by the NSF (Grants PHY-2019745, PHY-2014141, and PHY-2210291) and the Welch Foundation (Grant C-1792). ABOJ acknowledges the Robert A. Welch Postdoctoral Fellow Program. VGC acknowledges support from the NSF Award PHY-2609969 and the NVIDIA Academic Grant Program. JNO is a Cancer Prevention and Research Institute of Texas (CPRIT) Scholar in Cancer Research. We would like to thank AMD (Advanced Micro Devices, Inc.) for the donation of critical hardware and support resources from its HPC Fund that made this work possible.

## Appendix A: Experimental data analysis

The genome-scale imaging datasets from Ref. [8] were obtained from Ref. [38]. To assign the compartment annotation of each imaged locus, we used PyMEGABASE [39]to predict the subcompartment annotations for the human cell line IMR-90, employing all ChIP-seq, RNA-seq, and transcription factor tracks available for this cell line in the ENCODE database [40]. This annotation is available on the ENCODE portal under the accession code ENCFF174SZW. We then overlapped the predictions from PyMEGABASE with the genomic coordinates of the regions imaged.

Most of the analysis performed in our work involved structures from all five reported experiments [8]. We excluded any genome-wide structure that contains at least one chromosome whose asphericity deviated by more than three standard deviations from the mean asphericity of that chromosome across the ensemble.

Following the approach of Su et al. [8], we approximated the nuclear periphery by calculating the convex hull of all imaged loci for each cell in the ensemble. Here, we consider that a chromosome is in significant contact with the lamina (∈ lamina) if 10% or more of its imaged loci are on the boundary of the convex hull, i.e., points that have zero distance from the hull. In contrast, a chromosome is not in contact with the lamina (*∉* lamina) if none of its imaged loci is on the nuclear boundary.

We also used this convex hull of the nucleus to estimate the orientation of each chromosome relative to the nuclear periphery. For chromosomes with direct lamina contacts (loci with zero signed distance to the convex hull), we identified the nearest convex hull vertex for each contacting bead and assigned the corresponding vertex normal vector. We then define a chromosome lamina-normal vector, 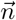, as the sum of the normals associated with all contacting A/B loci. For chromosomes without direct lamina contacts, we selected the 10% of chromatin beads closest to the convex hull (excluding unannotated beads) and similarly assigned each bead the normal vector of its nearest hull vertex. The chromosome lamina-normal vector was then obtained by summing these normals. This procedure yields an effective outward normal vector characterizing the local orientation of the nuclear surface adjacent to each chromosome.

To calculate the compartment polarization, 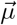, we used the PyMEGABASE annotations together with Equations 1 and 2. For each chromosome, the relative orientation between 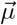 and 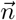 is calculated to be the angle defined by these two vectors, here called the *lamina-dipole angle*.

## Appendix B: Simulation Details

Chromosome dynamics simulations were performed using MiChroM [5] at a resolution of 50 kb. To match the cell line used in the experimental dataset [8], we used the same IMR-90 PyMEGABASE subcompartment an-notations as input for the model. The simulations were performed using the OpenMiChroM package [41].

### 1. Energetics of the model

Chromatin-chromatin interactions were modeled at the compartment level, using the following interaction parameters (Table I, previously described in Ref. [14]).

**TABLE I.**
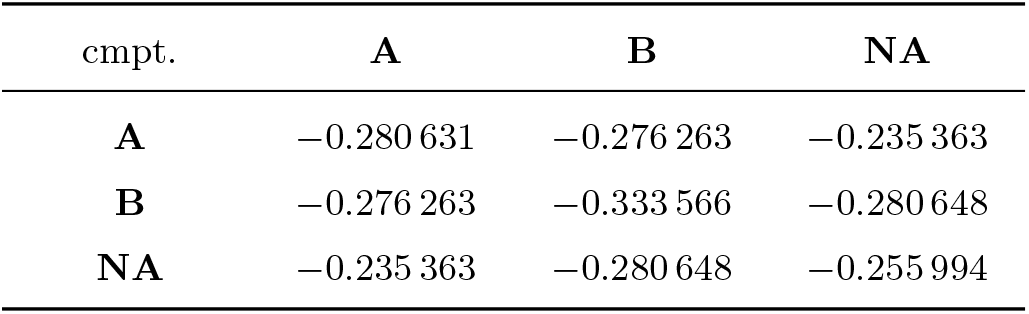
The parameters α_*kl*_ for chromatin-chromatin interactions, in units of ϵ.

In addition to chromatin-chromatin interactions, we introduced interactions between chromatin and the lamina and nucleoli. Following previous observations that each subcompartment interacts differently with each nuclear body [24, 42], we used the imaging experiments of Ref. [8] to estimate the interaction energies between subcompartments and nuclear bodies.

To estimate these values, we compared the frequency of contacts of a subcompartment *s* with a particular region *r* with the frequency of contact of a reference group of subcompartments. When *r* is the lamina, an imaged region belonging to the subcompartment *s* is defined in contact with *r* if it has zero distance to the nuclear convex hull (see Appendix A). When *r* represents the nucleoli, we use the distances to the nucleoli as reported in experiments 3 to 5 in Ref. [8], and a contact threshold of 0.10 *µ*m. We excluded chromosomes that contain rDNA regions (chromosomes 13, 14, 15, 21, and 22), as they are consistently found close to the nucleoli [8]. The different approaches follow the distance distributions extracted from the experimental data for each nuclear landmark [8] (Fig. S5). Therefore, in the simulation, the subcompartments chosen as reference subcompartments do not interact with that region, and the strength of interaction for the other subcompartments is defined by

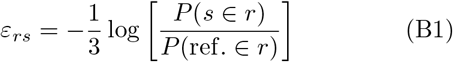

where *P*(*x* ∈*r*) represents the probability of *x* being in contact with *r* in the ensemble of structures. The factor of 1*/*3 compensates for the multiple chromatin-landmark interactions that result from representing the nuclear landmarks as beads. The computed values for the interaction strength are shown in Table II.

**TABLE II.**
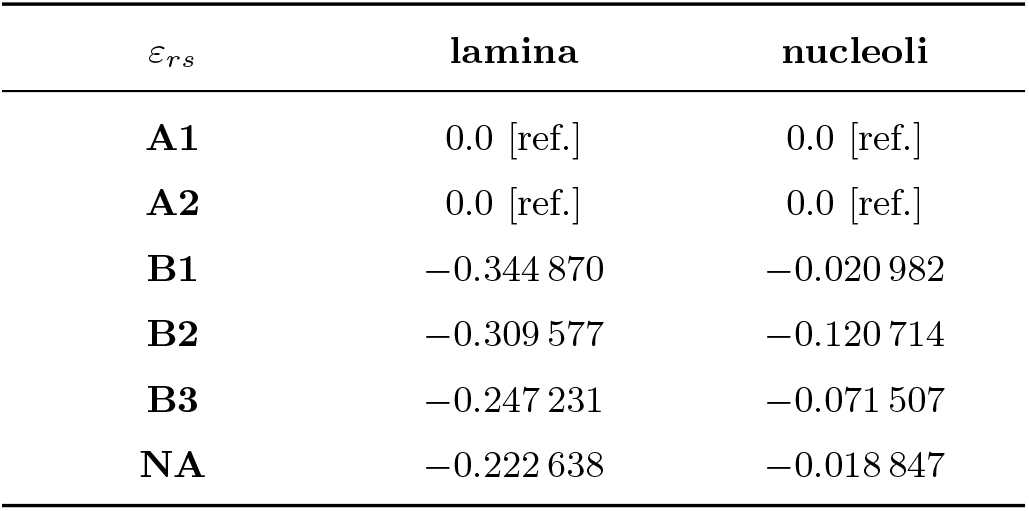
The parameters ε_*rs*_ for chromatin-landmark interactions, in units of ϵ.

The resulting Hamiltonian has the following form:

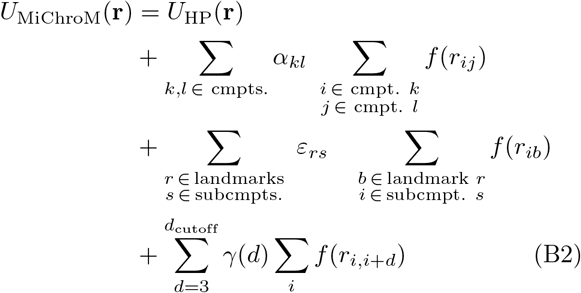

where *U*_HP_ is the homopolymer potential responsible for the polymeric nature of chromatin, including chain connectivity, bending stiffness, excluded volume, and permits chain crossing with an energetic penalty of 4 *k*_B_*T* . The second term represents the chromatin-chromatin interactions, modeled at the compartment level (Table I). The third term represents the chromatin-landmark interactions mentioned above (Table II). The last term in Eq. B2 is a lengthwise compaction that accounts for the characteristic decay of contact probability of chromosomes [2, 5, 13]. The function *f*(*r*) is a switch-like function that converts pairwise distances into contact probability, as follows:

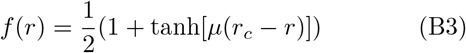

where *µ* = 3.22 *σ*^−1^ and *r*_*c*_ = 1.78 *σ*, with *σ* being the reduced unit of length.

### 2. Representation of nuclear landmarks

To determine the radius of the nucleus, we calculated the average volume for the imaged cells of Ref. [8], arriving at⟨*v*⟩ = 394.74 *µ*m^3^. We then calculated the radius of a sphere of this volume and converted it to reduced units using the relationship 1 *σ ≈* 0.14 *µ*m [16, 28], arriving at *R*_nucleus_ = 32.5 *σ*.

To model the nucleolus, we used a single nucleolus occupying 20% of the volume of the nucleus, i.e., with a radius of *R*_nucleolus_ = 19.0 *σ* [43]. Although the number of nucleoli per nucleus varies within a population of the same cell line and throughout the cell cycle, our goal here is not to capture that heterogeneity, but to isolate how contact with a nucleolar surface influences the structure of a chromosome territory [44, 45]. Therefore, a single static nucleolus centered in the nucleus provides the relevant surface for this interaction while keeping a minimal model, especially because chromosomes are simulated individually in the nuclear context.

The spherical confinement and the excluded volume of the nucleolus are modeled similarly to chromatin-chromatin excluded volume, but using the distance of chromatin beads to the respective spherical surface. To mimic the excluded volume of other chromosomes and control the volumetric density of the system, we applied a conic constraint aligned with the positive *z* axis. The half-opening angle *θ* of the cone was chosen such that the chromatin volume density was *ρ* = 0.30 [46], considering the total chromatin volume and the volume inside the cone and between the spheres (nucleus periphery and nucleolus). The conic confinement uses a flat-bottom harmonic potential based on the distance to the cone surface, with a force constant of *k* = 5× 10^−3^ *ϵ/σ*^2^, where *ϵ* is the reduced unit of energy.

Attractive interactions with nuclear landmarks were represented by immobile beads placed on the surfaces of the lamina and nucleolus. The beads were distributed on their respective spherical surfaces using a Fibonacci lattice on a sphere. The number of surface beads was chosen so that the mean nearest-neighbor spacing on the surface was approximately 1.0 *σ*, which corresponds to the diameter of the chromatin beads. For computational efficiency during sampling simulations, we only kept landmark beads that are within a conic region with a half-opening angle of *θ*_landmarks_ = 1.25 *θ*, where *θ* is the angle for the conic constraint.

### 3. Simulation setup

Using the described configuration, we simulated chromosome 1 of the human cell line IMR-90 with (∈ lamina) and without (*∉* lamina) the attractive interactions with the nuclear periphery. This is the only difference between the two sets of simulations, allowing us to isolate the effect of lamina interactions in the territory configuration. For each condition, we performed 96 independent trajectories. To obtain an initial collapsed structure of chr1 from PyMEGABASE annotations, we created a circular spring [41] and simulated the system for 5×10^3^ *τ* at an information temperature equal to 1.0, where *τ* is the reduced unit of time. We then performed two annealing cycles, raising the temperature to *T* = 2.0 over 1 ×10^4^ *τ*, cooling the system until *T* = 0.01 over an additional 2 ×10^4^ *τ*, bringing the temperature back to *T* = 1.0 over 1 *×*10^4^ *τ*, and repeating the procedure. We saved a minimized structure sampled at each *T* = 0.01 time point to be used as initial structures in the assembly of the nuclear context for each condition. The compact structure helps to position the chromatin between the nuclear periphery and the nucleolus.

This minimized chromosome structure was positioned inside the spherical nucleus. The structure was rotated so that the bead farthest from the center was aligned with the positive *z* axis. The chromosome was then translated so that the gap between the chromosome and the nuclear periphery was 1.5 *σ*. The nucleolus was initially positioned along the *z* axis below the lowest *z* coordinate of the chromosome, also with a gap of 1.5 *σ*. A short simulation was performed to gradually move the nucleolus until it was positioned in the center of the system. Following an equilibration of 5 ×10^3^ *τ* at *T* = 1.0, the nucleolus was moved over 100 intervals of 50 *τ* . After the nucleolus reached the origin, the system was annealed by increasing the temperature from *T* = 1.1 to *T* = 2.0 over 10^4^ *τ* and then cooling from *T* = 1.99 to *T* = 1.0 in another 10^4^ *τ* . A final equilibration of 5 ×10^3^ *τ* was performed before saving the assembled state. The intent of this assembly procedure was to avoid large initial overlaps between the chromosome and the excluded volume from the nuclear periphery and the nucleolus.

### 4. Equilibration and sampling

For production sampling, the assembled state was reloaded, adding the conic confinement and filtering the explicit landmark beads for each condition (see Appendix B2). The same annealing schedule was then repeated: an initial equilibration of 5 ×10^3^ *τ*, heating to *T* = 2.0 over 10^4^ *τ*, cooling to *T* = 1.0 over 10^4^ *τ*, and a final equilibration of 5 ×10^3^ *τ* . Sampling was then performed for 10^5^ *τ*, recording structures every 10 *τ* . With 96 independent trajectories for each condition, we collected a total of 960,000 structures per condition.

