## Supporting Information for "The compartmental polarization of interphase human chromosomes within the nucleus: A mechanism for transcriptional regulation of large chromosomes"

Matheus F. Mello
*Center for Theoretical Biological Physics, Rice University, Houston, TX 77005 and Systems, Synthetic, and Physical Biology PhD Program, Rice University, Houston, TX 77005*

Dimanthi N. Perera
*Center for Theoretical Biological Physics, Rice University, Houston, TX 77005 and Department of Biology and Biochemistry, University of Houston, Houston, TX 77004*

Lauren E. DiPaola
*Center for Theoretical Biological Physics, Rice University, Houston, TX 77005 and Department of Biomedical Engineering, University of Houston, Houston, TX 77004*

Antonio B. Oliveira Jr.
*Center for Theoretical Biological Physics, Rice University, Houston, TX 77005 and Department of Physics, São Paulo State University (UNESP), Institute of Biosciences, Humanities and Exact Sciences, São José do Rio Preto, São Paulo 15054-000, Brazi*

Vinícius G. Contessoto
*Center for Theoretical Biological Physics, Rice University, Houston, TX 77005 and Department of Physics and Astronomy, Rice University, Houston, TX 77005*

José N. Onuchic
*Center for Theoretical Biological Physics, Rice University, Houston, TX 77005 Department of Physics and Astronomy, Rice University, Houston, TX 77005 Department of Chemistry, Rice University, Houston, TX 77005 and Department of Biosciences, Rice University, Houston, TX 77005*

Ryan R. Cheng
*Department of Chemistry, University of Kentucky, Lexington, KY 40506* *

(Dated: August 21, 2026)

### I. EXTENDED METHODS

#### A. MiChroM potential

The MiChroM potential is derived from the Maximum Entropy Approach [1, 2] and proposes a energy function for chromatin based on the phase-separation of active and inactive chromatin, and on the lengthwise compaction resulting from SMC motors activity. On top of the original model, that here is used at the compartment level, i.e., each bead represents a 50 kb locus that belongs to compartment A (active chromatin), B (inactive chromatin), or NA (non annotated), we have included chromatin interactions with the lamina and the nucleolus. The resulting potential has the following form

$$U_{\mathrm{MiChroM}}(\mathbf{r}) = U_{\mathrm{HP}}(\mathbf{r}) + \underbrace{\sum_{\substack{k \geq l \\ k,l \,\in\, \text{cmpts.}}} \alpha_{kl} \sum_{\substack{i \,\in\, \text{cmpt. } k \\ j \,\in\, \text{cmpt. } l}} f(r_{ij})}_{\text{chromatin-chromatin interactions}}$$

$$+ \underbrace{\sum_{\substack{r \,\in\, \text{landmarks} \\ s \,\in\, \text{subcmpts.}}} \varepsilon_{rs} \sum_{\substack{b \,\in\, \text{landmark } r \\ i \,\in\, \text{subcmpt. } s}} f(r_{ib})}_{\text{chromatin-landmark interactions}} + \underbrace{\sum_{d=3}^{d_{\text{cutoff}}} \gamma(d) \sum_{i} f(r_{i,i+d})}_{\text{ideal chromosome}} \tag{S1}$$

where $U_{\mathrm{HP}}(\mathbf{r})$ accounts for the energetic contribution of the polymer, including the potentials for bonds, angles, and excluded volume. The parameters $\alpha_{kl}$ represent the strength of interaction between loci of compartments $k$ and $l$, respectively. The function $f(r_{ij})$ is a switch-like function that converts Euclidean distance into contact probability. The parameters $\varepsilon_{rs}$ represent the strength of interaction between a chromatin bead and a nuclear-landmark bead, here the lamina or nucleolus. Finally, the parameters in $\gamma(d)$ depend on the genomic distance along the chain $d$ and control the lengthwise compaction of the polymer.

The contact switch function has the form of

$$f(r) = \frac{1}{2}\left(1 + \tanh\left[\mu(r_c - r)\right]\right) \tag{S2}$$

where the parameters $\mu$ and $r_c$ control the shape of the function. To determine these parameters, two beads are considered in contact, i.e., $f = 1$, if $r_{ij} = 1\ \sigma$, since the diameter of the bead is $1\ \sigma$. The second constraint is the contact probability extracted from the experimental Hi-C map for the second neighbor, resulting in

$$f(1) = 1 \qquad f(2) = \min\left\{P^{\exp}_{i,i+2}\right\}$$

where we assume $\bar{r}_{i,i+2} = 2\ \sigma$ since the equilibrium angle for the angle potential is considered to be $\theta_0 = \pi$ rad, as described further in the text. The resulting parameters calculated in Ref. [2] for the human cell line GM12878 were $\mu = 3.22\ \sigma^{-1}$ and $r_c = 1.78\ \sigma$, and were shown to be transferable to other cell lines [3, 4].

#### B. Homopolymer model

The homopolymer model used in MiChroM represents a basic polymeric model in which each bead corresponds to a genomic locus of 50 kb. It can be expressed as follows

$$U_{\mathrm{HP}}(\mathbf{r}) = \sum_{i \,\in\, \{\text{chr. loci}\}} \left[ U_{\mathrm{FENE}}(r_{i,i+1}) + U_{\mathrm{hc}}(r_{i,i+1}) + U_{\mathrm{periphery}}(r_i) + U_{\mathrm{nucleolus}}(r_i) + U_{\mathrm{conic}}(r_i) \right]$$

$$+ \sum_{i \,\in\, \{\text{angles}\}} U_{\mathrm{angle}}(\theta_i) + \sum_{\substack{j \,>\, i+2 \\ i,j \,\in\, \{\text{chr. loci}\}}} U_{\mathrm{sc}}(r_{i,j}) \tag{S3}$$

$U_{\mathrm{FENE}}$ represents the Finite Extensible Nonlinear Elastic (FENE) bonding potential, which is applied to bonded monomers and has the following form:

$$U_{\mathrm{FENE}}(r) = \begin{cases} -\dfrac{1}{2} k_b R_0^2 \ln\left[1 - \left(\dfrac{r}{R_0}\right)\right] & \text{for} \quad r \leq R_0 \\ 0 & \text{for} \quad r > R_0 \end{cases}$$

A hard-core repulsive potential is also added to bonded monomers to avoid overlap.

$$U_{\mathrm{hc}}(r) = \begin{cases} 4\varepsilon \left[\left(\dfrac{\sigma}{r}\right)^{12} - \left(\dfrac{\sigma}{r}\right)^{6} + \dfrac{1}{4}\right] & \text{for} \quad r \leq \sigma 2^{\frac{1}{6}} \\ 0 & \text{for} \quad r > \sigma 2^{\frac{1}{6}} \end{cases}$$

$U_{\text{angle}}$ represents the angular potential applied to all three consecutive beads and assumes the form

$$U_{\text{angle}}(\theta_i) = k_a\left[1 - \cos(\theta_i - \theta_0)\right] \tag{S4}$$

where $\theta_i$ is the angle between the vectors $\vec{r}_{i,i+1}$ and $\vec{r}_{i,i-1}$ and $\theta_0$ is the resting angle.

A soft-core repulsive interaction is added between non-bonded pairs:

$$U_{sc}(r) = \frac{1}{2}E_{cut}\left[1 + \tanh\left(1 - k_{sc}(r - r_0)\right)\right] \tag{S5}$$

where $k_{sc} = 21.94$ and $r_0 = 0.93\ \sigma$. Here we propose an alternative functional form to the soft-core potential originally described in Ref. [2], which uses the Lennard-Jones potential in its definition. However, given the relatively large time step we use in our coarse-grained simulations, the Lennard-Jones potential can become numerically unstable when performing the computations on GPUs. Therefore, the values of $k_{sc}$ and $r_0$ were optimized such that the functional form of our proposed soft-core potential was consistent with that previously proposed [2].

Finally, the confinement potentials $U_{\text{periphery}}$ and $U_{\text{nucleolus}}$ represent the excluded volume of the nuclear periphery and the nucleolus, respectively.

$$U_{\text{periphery}}(r) = 5\,U_{sc}(R_{\text{nucleus}} - r) \tag{S6}$$

$$U_{\text{nucleolus}}(r) = 5\,U_{sc}(r - R_{\text{nucleolus}}) \tag{S7}$$

where $R_{\text{nucleus}}$ and $R_{\text{nucleolus}}$ are the radius of the nucleus and nucleolus, respectively. The factor of 5 ensures that the soft-core potential effectively behaves as hard-core potential while still being numerically stable. Note that since we use $U_{sc}$ in the definition of the landmark potentials, the lamina and nucleolus have a thickness of $0.5\ \sigma$, in alignment with the beads with a unitary diameter distributed on the surface of these landmarks used for attractive interactions with chromatin.

To mimic the excluded volume of other chromosomes in the nucleus, a conic confinement $U_{\text{conic}}$ is used. The vertex of the cone is at the origin of the system, and the conic potential is aligned with the positive $z$ axis.

$$U_{\text{conic}}(r) = \begin{cases} \frac{1}{2}k_{\text{cone}}\,d_{\text{cone}}^2 & \text{for } d_{\text{cone}} \geq 0 \\ 0 & \text{for } d_{\text{cone}} < 0 \end{cases} \tag{S8}$$

where $d_{\text{cone}} = \sqrt{x^2 + y^2}\,\cos(\phi) - z\,\sin(\phi)$, with $\phi$ is the half-opening angle of the cone.

The following values of parameters were used

$$k_a = 2\ \epsilon \qquad k_b = 30\ \frac{\epsilon}{\sigma^2} \qquad k_{\text{cone}} = 5\times10^{-3}\ \frac{\epsilon}{\sigma^2} \qquad E_{\text{cut}} = 4\ \epsilon \qquad R_0 = 1.5\ \sigma \qquad \theta_0 = \pi\ \text{rad}$$

where $\epsilon = k_{\text{B}}T$ is the reduced unit of energy, and $\sigma$ is the reduced unit of length, with the bead diameter being $1\ \sigma$.

#### C. MiChroM parameter set

The chromatin-chromatin interactions are determined by the parameters $\alpha_{kl}$, which set the strength of interaction between two beads of compartments $k$ and $l$. The parameters for MiChroM at the compartment level were first determined by Ref. [3] and are defined in Table S1.

The set of parameters of the ideal chromosome potential is defined by

$$\gamma(d) = \frac{\gamma_1}{\log(d)} + \frac{\gamma_2}{d} + \frac{\gamma_3}{d^2} \tag{S9}$$

where $d$ is the genomic distance and $\gamma_1 = -0.030\ \epsilon$, $\gamma_2 = -0.351\ \epsilon$ and $\gamma_3 = -3.727\ \epsilon$. The ideal chromosome potential was applied until $d_{\text{cutoff}} = 500$, i.e., between beads $i$ and $j$ if $3 \leq |i - j| \leq 500$.

To estimate the parameters for the chromatin-landmark interactions, we used the imaging experiments of Ref. [5]. More specifically, we compared the frequency of contacts of a subcompartment $s$ with a particular region $r$ with the frequency of contact of a reference group of subcompartments. When $r$ is the lamina, an imaged region belonging to

TABLE S1. The parameters $\alpha_{kl}$ for chromatin-chromatin interactions, in units of $\epsilon$.

| cmpt. | **A** | **B** | **NA** |
|---|---|---|---|
| **A** | −0.280 631 | −0.276 263 | −0.235 363 |
| **B** | −0.276 263 | −0.333 566 | −0.280 648 |
| **NA** | −0.235 363 | −0.280 648 | −0.255 994 |

TABLE S2. The parameters $\varepsilon_{rs}$ for chromatin-landmark interactions, in units of $\epsilon$.

| $\varepsilon_{rs}$ | **lamina** | **nucleoli** |
|---|---|---|
| **A1** | 0.0 [ref.] | 0.0 [ref.] |
| **A2** | 0.0 [ref.] | 0.0 [ref.] |
| **B1** | −0.344 870 | −0.020 982 |
| **B2** | −0.309 577 | −0.120 714 |
| **B3** | −0.247 231 | −0.071 507 |
| **NA** | −0.222 638 | −0.018 847 |

the subcompartment $s$ is defined in contact with $r$ if it has zero distance to the nuclear convex hull (see Appendix A). When $r$ represents the nucleoli, we use the distances to the nucleoli as reported in experiments 3 to 5 in Ref. [5], and a contact threshold of 0.10 $\mu$m. We excluded chromosomes that contain rDNA regions (chromosomes 13, 14, 15, 21, and 22), as they are consistently found close to the nucleoli [5]. The different approaches follow the distance distributions extracted from the experimental data for each nuclear landmark [5] (Fig. S5).

Therefore, in the simulation, the subcompartments chosen as reference subcompartments do not interact with that region, and the strength of interaction for the other subcompartments is defined by

$$\varepsilon_{rs} = -\frac{1}{3} \log \left[ \frac{P(s \in r)}{P(\text{ref.} \in r)} \right] \tag{S10}$$

where $P(x \in r)$ represents the probability of $x$ being in contact with $r$ in the ensemble of structures. The factor of $1/3$ compensates for the multiple chromatin-landmark interactions that result from representing the nuclear landmarks as beads. The computed values for the interaction strength are shown in Table S2.

#### D. Simulation details

Chromosome simulations were performed using the Minimal Chromatin Model (MiChroM) [2] through Open-MiChroM [6], with the modifications mentioned above. To match the experimental genome-wide imaging data from Ref. [5], all simulations used the IMR-90 subcompartment annotation predicted by PyMEGABASE [7], available under the ENCODE accession number `ENCFF174SZW`. Simulations are performed for chromosome 1 of this cell line at a resolution of 50 kb, with a time step $\Delta t = 0.01\tau$, where $\tau$ is the reduced time unit of the model.

The nucleus was modeled as a sphere centered at the origin of the system. To determine the radius of the nucleus, we calculated the average volume for the imaged cells of Ref. [5], arriving at $\langle v \rangle = 394.74\ \mu\text{m}^3$. We then calculated the radius of a sphere of this volume and converted it to reduced units using the relationship $1\ \sigma \approx 0.14\ \mu$m [8, 9], arriving at $R_{\text{nucleus}} = 32.5\ \sigma$. The nucleolus was modeled as a single sphere placed at the origin that occupies 20% of the volume of the nucleus, i.e., with a radius of $R_{\text{nucleolus}} = 19.0\ \sigma$ [10]. The half-opening angle $\phi$ of the conic confinement is calculated such that the volume density of the chromatin in the volume inside the cone and between the two spheres of the nucleus and the nucleolus is 0.30 [11].

Our goal is to investigate the effects of lamina interactions on the structure of the chromosome territory; therefore, we performed two groups of simulations, with ($\in$ lamina) and without ($\notin$ lamina) attractive interactions between the chromatin beads and the lamina. The simulations for both conditions shared the same simulation protocol, and they differ only by the presence of attractive interactions with the lamina. The simulation protocol is divided into three parts: initial collapse, assembly, and sampling, and each of them is described below.

*1. Initial collapse*

To obtain an initial collapsed structure of the chromosome, we created a circular spring [6] and submitted it to a flat-bottom harmonic potential $U_{\text{collapse}}$ to aid the collapse. The potential has the form

$$U_{\text{collapse}}(r_i) = \begin{cases} k_r(r_i - R_r)^2 & \text{for} \quad r_i \geq R_r \\ 0 & \text{for} \quad r_i < R_r \end{cases} \tag{S11}$$

where $k_r = 5 \times 10^{-3} \ \frac{\epsilon}{\sigma^2}$ and $R_r = 10 \ \sigma$. The system was simulated for $5 \times 10^3 \ \tau$ at an information temperature equal to 1.0, after which annealing cycles were performed. The temperature was raised from $T = 1.0$ to $T = 2.0$ in 100 increments of $10^2 \ \tau$, cooled to $T = 0.01$ in 200 increments of $10^2 \ \tau$, and brought back to $T = 1.0$ in another 100 increments of $10^2 \ \tau$. The procedure is repeated once more and, at each $T = 0.01$ time point, a annealed structure is sampled to be used as initial structures in the assembly of the nuclear context for each simulated conditions. This compact structure helps to position the chromatin between the nuclear periphery and the nucleolus.

*2. Assembly*

The assembly phase consists of positioning the chromosome within the nuclear context. The annealed chromosome structure is loaded inside the spherical nucleus. The structure is then rotated so that the bead farthest from the center aligns with the positive $z$ axis. To construct the lamina and nucleolus beads to be used in the attractive interactions with the nuclear landmarks, we iteratively generated a Fibonacci lattice on a sphere with the appropriate radius until the mean nearest-neighbor spacing on the surface was approximately 1.0 $\sigma$. These beads are immobile during the simulation and act as fixed interaction sites. The chromosome is then translated so that the gap between the chromosome and the nuclear periphery is 1.5 $\sigma$. The nucleolus is initially positioned along the $z$ axis below the lowest $z$ coordinate of the chromosome, also with a gap of 1.5 $\sigma$.

A short simulation is performed to gradually move the nucleolus until it is positioned in the center of the system. Following an equilibration of $5 \times 10^3 \ \tau$ at $T = 1.0$, the nucleolus is moved over 100 intervals of 50 $\tau$. After the nucleolus reached the origin, the system is annealed again by increasing the temperature from $T = 1.0$ to $T = 2.0$ in 100 increments of $10^2 \ \tau$ and then cooling from $T = 1.99$ to $T = 1.0$ in another 100 increments of $10^2 \ \tau$. A final equilibration of $5 \times 10^3 \ \tau$ is performed before saving the assembled state. The purpose of this assembly procedure is to avoid large initial overlaps between the chromosome and the excluded volume of the nuclear periphery and the nucleolus.

*3. Sampling*

For production sampling, the chromosome from the assembled state is loaded and re-aligned with the $z$ axis. The conic confinement is added to the system. For computational efficiency, we remove from the system the landmark beads that lie outside a conic region with a half-opening angle of 1.25 $\phi$, where $\phi$ is the half-opening angle for the conic constraint. The same annealing schedule is then repeated: an initial equilibration of $5 \times 10^3 \ \tau$, heating to $T = 2.0$ in 100 increments of $10^2 \ \tau$, cooling to $T = 1.0$ in 100 increments of $10^2 \ \tau$, and a final equilibration of $5 \times 10^3 \ \tau$. Then, sampling is performed for $10^5 \ \tau$, recording the coordinates of the system every 10 $\tau$. For each condition, 96 independent trajectories were performed, totaling 960,000 sampled structures per condition.

### II. DATA ANALYSIS

#### A. Analysis of experimental imaging data

The genome-scale imaging datasets from Ref. [5] were obtained from Ref. [12]. To assign the compartment annotation of each imaged locus, we used PyMEGABASE [7] to predict the subcompartment annotations for the human cell line IMR-90, employing all ChIP-seq, RNA-seq, and transcription factor tracks available for this cell line in the ENCODE database [13]. This annotation is available on the ENCODE portal under the accession code `ENCFF174SZW`. We then overlapped the predictions from PyMEGABASE with the genomic coordinates of the regions imaged.

Most of the analysis performed in our work involved structures from all five reported experiments [5]. We excluded any genome-wide structure that contains at least one chromosome whose asphericity deviated by more than three standard deviations from the mean asphericity of that chromosome across the ensemble.

Following the approach of Su et al. [5], we approximate the nuclear periphery by calculating the convex hull of all imaged loci for each cell in the ensemble. Here, we consider that a chromosome is in significant contact with the lamina ($\in$ lamina) if 10% or more of its imaged loci are on the boundary of the convex hull, i.e., points that have zero distance from the hull. In contrast, a chromosome is not in contact with the lamina ($\notin$ lamina) if none of its imaged loci is on the nuclear boundary.

#### B. Compartment polarization

For both experimental structures and simulated trajectories, the compartment polarization was calculated as

$$\vec{\mu} = \sum_i q_i(\vec{r}_i - \vec{r}_{\mathrm{COM}}), \tag{S12}$$

where $\vec{r}_i$ is the position of bead or locus $i$, $\vec{r}_{\mathrm{COM}}$ is the center of mass of the chromosome, and

$$q_i = \begin{cases} +1, & i \in \text{A compartment} \\ -1, & i \in \text{B compartment} \\ 0, & i \in \text{NA} \end{cases}. \tag{S13}$$

To calculate the degree of alignment of the compartment dipole with the lamina, we first need to define a proxy for the normal direction to the lamina. For the experimental structures, we used the nuclear convex hull to estimate the orientation of each chromosome relative to the nuclear periphery. For chromosomes with direct lamina contacts (loci with zero signed distance to the convex hull), we identified the nearest convex hull vertex for each contacting bead and assigned the corresponding vertex normal vector. We then define a chromosome lamina-normal vector, $\vec{n}$, as the sum of the normals associated with all A/B contacting loci. For chromosomes without direct lamina contacts, we selected the 10% of chromatin beads closest to the convex hull (excluding unannotated beads) and similarly assigned each bead the normal vector of its closest hull vertex. The chromosome lamina-normal vector was then obtained by summing these normals. This procedure yields an effective outward normal vector characterizing the local orientation of the nuclear surface adjacent to each chromosome. In the simulated trajectories, we approximated the lamina-normal vector $\vec{n}$ to have the same direction of the vector defined by the nuclear center (system's origin) to the chromosome center of mass.

The lamina-dipole alignment angle is then calculated as

$$\theta = \arccos\left[\frac{\vec{\mu}\cdot\vec{n}}{|\vec{\mu}||\vec{n}|}\right]. \tag{S14}$$

#### C. Contact analysis

When counting the number of contacts, a binary contact definition was used. Two loci $i$ and $j$ were considered in contact if $f(r_{ij}) \geq 0.5$, which corresponds to $r_{ij} \leq r_c = 1.78\ \sigma$. Contacts were not considered for first and second neighbors.

To investigate the dynamics of contact formation, we calculated, for each bead $i$ in compartment $C$, the number of contacts $N_i^C(t)$ it makes at time $t$ with other beads $j$ in the same compartment, where $C \in \{\mathrm{A}, \mathrm{B}, \mathrm{NA}\}$ and $|i-j| \geq 3$. We defined $t_{\mathrm{on}}$ as the duration of a continuous time interval during which $N_i^C(t) > 0$, indicating that the bead $i$ maintains at least one same-compartment contact. In contrast, $t_{\mathrm{off}}$ was defined as the duration of a continuous time interval during which $N_i^C(t) = 0$, indicating that the bead $i$ has no contact with beads in the same compartment.

#### D. Chromatin Density Profile

To characterize the spatial localization of chromatin compartments within the simulated territory, we leverage the cylindrical symmetry of the simulation setup and calculated the density profile in the system. For that, in each frame the chromosome coordinate was rotated in relation to the origin of the system so that the vector from the center to the chromosome center of mass was aligned with the positive $z$ axis. After representing the chromatin positions in spherical coordinates $(r, \theta, \phi)$, we calculated the number of beads as a function of $(r, \theta)$, integrating out the azimuthal coordinate $\phi$. The final normalized density $\rho_C(r, \theta)$ for compartment $C \in \{\mathrm{A}, \mathrm{B}\}$ can be defined as

$$\rho(r, \theta) = \frac{\langle N_C(r, \theta) \rangle}{N_{C,\,\mathrm{total}} \cdot \Delta V(r, \theta)} \tag{S15}$$

where $N_{C,\,\mathrm{total}} = \sum_{r,\theta} N_C(r, \theta)$ represents the total number of loci belonging to compartment $C$ and

$$\Delta V(r, \theta) = \frac{2\pi}{3} \left( (r + \Delta r)^3 - r^3 \right) [\cos(\theta) - \cos(\theta + \Delta\theta)] \tag{S16}$$

We used 10 linear radial bins between $R_{\mathrm{nucleolus}} + 0.5\ \sigma$ and $R_{\mathrm{nucleus}} - 0.5\ \sigma$, and 20 linear angular bins between 0 and $\pi/4$. Thus, the density profile maps of each condition represent the probability of finding an A- or B-compartment bead in each spatial region, normalized by the total number of A or B loci and the region's volume.

### E. Cluster analysis

To quantify the formation of heterochromatin-rich clusters, we analyzed the connectivity of B-compartment beads. Using the same contact threshold ($r_{ij} \leq r_c = 1.78\ \sigma$ and $|i - j| \geq 3$), for each simulated frame, we constructed an undirected contact graph in which each B-compartment bead was represented by a node and an edge was assigned between every pair of beads that met the contact criterion a graph of the beads connected by at least one contact, excluding first and second neighbor-contacts. The group of connected beads are then defined as a cluster. Isolated beads were treated as clusters of size one.

For each structure, we calculated the fraction of B loci that participate in the largest cluster,

$$f_{\max}(t) = \frac{N_{\max}(t)}{N_{\mathrm{B}}}, \tag{S17}$$

where $N_{\max}(t)$ is the number of beads in the largest connected B-compartment cluster at time $t$, and $N_{\mathrm{B}}$ is the total number of B-compartment beads in the chromosome. We calculated $f_{\max}$ for all sampled frames of each independent trajectory.

### F. Software and computational tools

All structural representations shown in this work were generated in VMD [14] and rendered with Tachyon [15]. Simulated trajectories were processed using the `CNDBTools` module distributed with OpenMiChroM [6]. Mean-squared displacements were calculated with `freud` [16]. Convex hull calculations used the `trimesh` module, available in `https://trimesh.org`. Additional codes used during the analysis of our trajectories can be found in the `chroma` package, available at `https://github.com/mellofariam/chroma`. The simulation and analysis scripts can also be found in `https://github.com/mellofariam/polarization`.

### III. SUPPLEMENTARY TABLES

TABLE S3. Moments $\langle X^n \rangle$ of the contact time distributions. The moments are computed for each simulated trajectory, and values are reported as $\langle\langle X^n \rangle\rangle_{\text{trajectories}} \pm \sigma(\langle X^n \rangle)$, in units of $\tau^n$.

| | | A | | B | |
|---|---|---|---|---|---|
| | $n$ | $\in$ lamina | $\notin$ lamina | $\in$ lamina | $\notin$ lamina |
| $\langle {t_{\text{on}}}^n \rangle$ | 1 | **3.98 ± 0.03 × $10^2$** | 2.88 ± 0.03 × $10^2$ | 4.91 ± 0.06 × $10^2$ | 4.82 ± 0.05 × $10^2$ |
| | 2 | **7.08 ± 0.56 × $10^4$** | 2.79 ± 0.21 × $10^4$ | **4.39 ± 0.47 × $10^5$** | 2.92 ± 0.23 × $10^5$ |
| | 3 | **6.41 ± 0.66 × $10^1$** | 5.12 ± 0.55 × $10^1$ | **1.59 ± 0.14 × $10^2$** | 1.20 ± 0.10 × $10^2$ |
| | 4 | **7.80 ± 1.90 × $10^2$** | 4.92 ± 1.25 × $10^2$ | **4.02 ± 0.67 × $10^3$** | 2.42 ± 0.42 × $10^3$ |
| $\langle {t_{\text{off}}}^n \rangle$ | 1 | 1.37 ± 0.01 × $10^1$ | 1.36 ± 0.01 × $10^1$ | 1.52 ± 0.01 × $10^1$ | 1.51 ± 0.01 × $10^1$ |
| | 2 | 1.25 ± 0.06 × $10^1$ | 1.16 ± 0.08 × $10^1$ | 2.44 ± 0.35 × $10^1$ | 2.13 ± 0.15 × $10^1$ |
| | 3 | 8.94 ± 1.02 × $10^1$ | 9.02 ± 1.11 × $10^1$ | **2.22 ± 1.52 × $10^2$** | 1.43 ± 0.45 × $10^2$ |
| | 4 | 1.90 ± 0.78 × $10^3$ | 2.08 ± 0.79 × $10^3$ | **2.41 ± 3.94 × $10^4$** | 8.23 ± 8.28 × $10^3$ |

### IV. SUPPLEMENTARY FIGURES

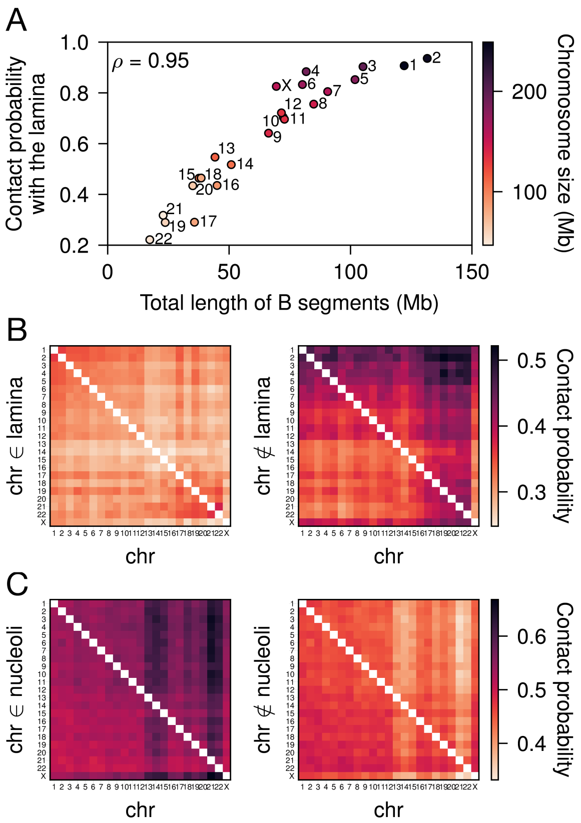


FIG. S1. **Chromosome interactions with nuclear landmarks and their effects in chromosome neighborhood**. (**A**) For genome-scale images of IMR-90 by Ref. [5], the contact probability of each chromosome with the nuclear lamina correlates with the total length of B segments, with a Spearman's correlation of $\rho = 0.95$. The compartment annotation for each chromosome was determined by MEGABASE for IMR-90 [7]. The lamina is estimated by calculating the convex hull all imaged loci in a cell [5]. Here, a chromosome is considered in contact with the lamina if any of its locus define the convex hull. The two homologs of each chromosome are averaged together. (**B**) Conditional contact probability among chromosomes, when the chromosome on the y-axis is in contact (left) or not in contact (right) with the lamina. Contact with the lamina is defined as in (A). (**C**) Conditional contact probability among chromosomes, when the chromosome on the y-axis is in contact (left) or not in contact (right) with the nucleoli. Chromosomes are considered in contact with the nucleoli when the closest distance to a nucleolus is $\leq 0.10$ $\mu$m. For both (B) and (C), inter-chromosomal contacts were identified in each imaged structure by computing contacts from Voronoi cells generated from each imaged locus. Chromosomes were considered in contact if they had at least one Voronoi contact for which the inter-loci distance was less than 1 $\mu$m.

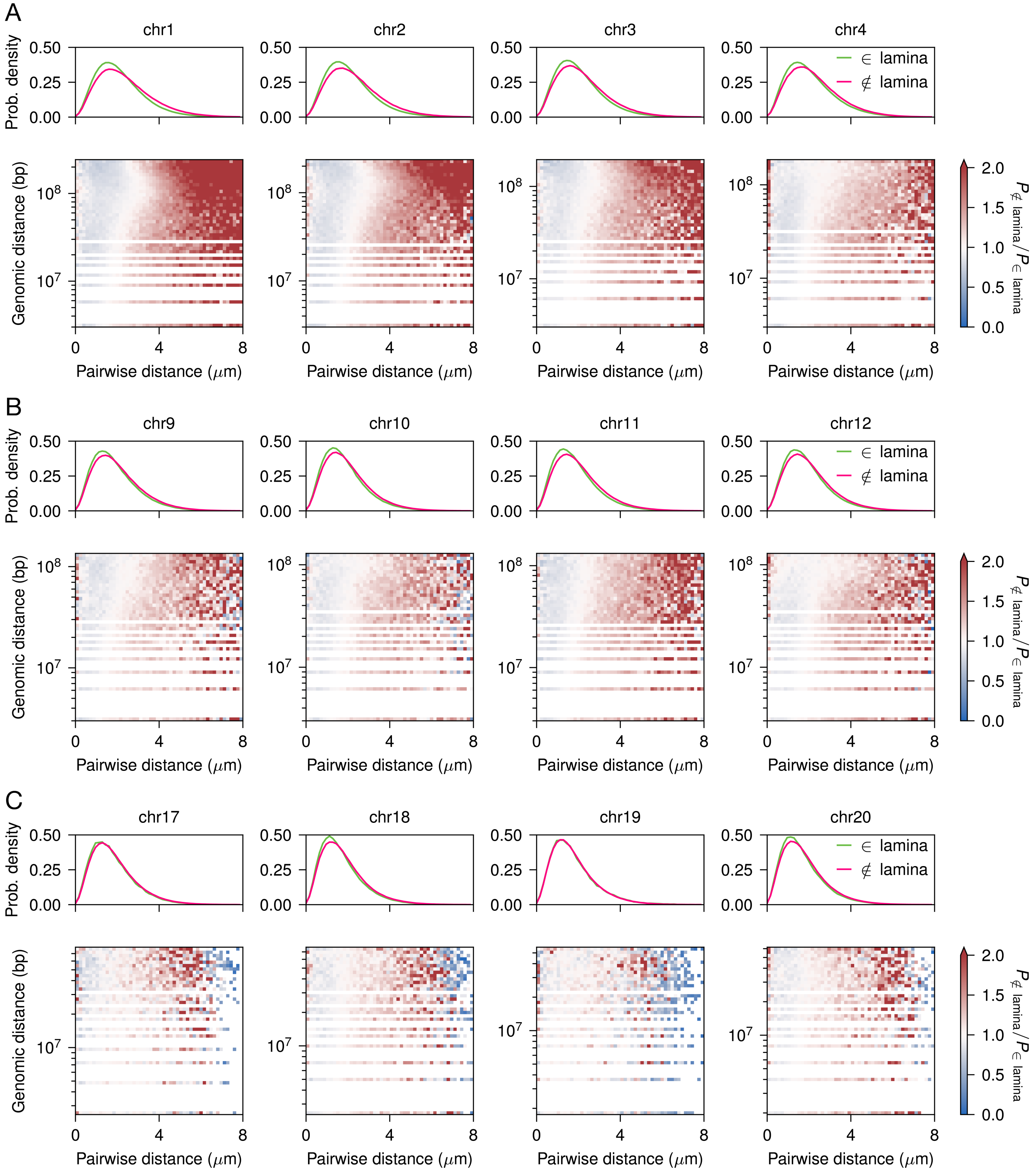


FIG. S2. **Comparison between the internal distances of chromosomes that interact or not with the nuclear periphery**. For the homologs of selected chromosomes in the whole-genome ensemble imaged by Ref. [5], pairwise distance distributions when the chromosome interacts with the lamina or not (top) and log ratio between the bi-variate probability density distributions $P_{\notin \text{ lamina}}(d_{ij}, r_{ij})/P_{\in \text{ lamina}}(d_{ij}, r_{ij})$ (bottom), where $d_{ij}$ and $r_{ij}$ respectively represent the genomic and spatial distances between loci $i$ and $j$. Four chromosomes were selected among large chromosomes (**A**), medium-sized chromosomes (**B**), and small chromosomes (**C**).

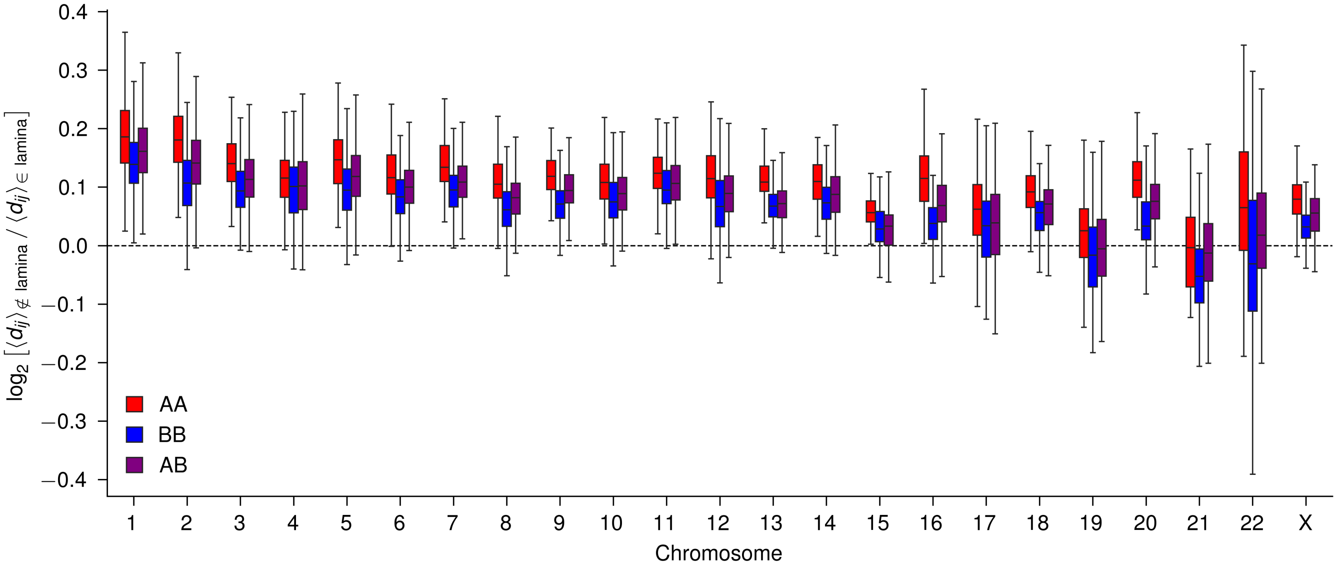


FIG. S3. **Differences in average pairwise imaged distances between the $\in$ lamina and $\notin$ lamina conditions are more prominent for euchromatin pairs in large chromosomes.**. For each chromosome in the whole-genome ensemble imaged by Ref. [5], the $\log_2$ ratio of the average pairwise distance is shown for each type of pair (AA, BB, or AB).

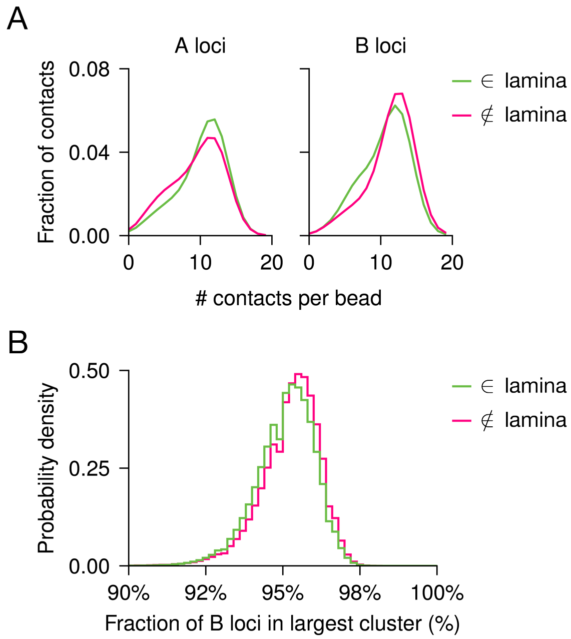


FIG. S4. **Shifts in the localization of compartments in the simulated ensemble of chr1 have no significant effects on the heterochromatin contact network**. (**A**) Histogram of the total number of contacts for A (left) and B (right) loci in the IMR-90 simulations of chr1. A pair of loci $(i, j)$ is considered in contact at a certain time $t$ in the simulation if $d_{ij}(t) \le r_c$, with $r_c = 1.78\ \sigma$ defined in Eq. S2. (**B**) Fraction of B loci in the largest B-loci cluster for the $\in$ and $\notin$ lamina conditions. For each simulated conformation, B loci were represented as a contact graph, with edges connecting non-neighboring B loci whose spatial separation satisfied the same contact threshold used in (A). The cluster size was computed as the fraction of all B loci belonging to the largest connected component of this graph.

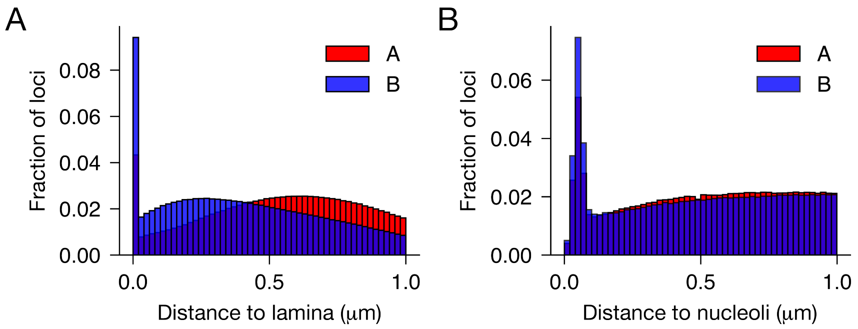


FIG. S5. **Imaged distances to nuclear landmarks**. (**A**) Distance to the inferred lamina for the whole-genome ensemble imaged by Ref. [5]. (**B**) Distance to the nucleoli as reported in experiments 3, 4, and 5 of whole-genome images by Ref. [5].

[1] E. T. Jaynes, Physical Review **106**, 620 (1957).
[2] M. D. Pierro, B. Zhang, E. L. Aiden, P. G. Wolynes, and J. N. Onuchic, Proceedings of the National Academy of Sciences **113**, 12168 (2016).
[3] M. D. Pierro, R. R. Cheng, E. L. Aiden, P. G. Wolynes, and J. N. Onuchic, Proceedings of the National Academy of Sciences **114**, 12126 (2017).
[4] R. R. Cheng, V. G. Contessoto, E. Lieberman Aiden, P. G. Wolynes, M. Di Pierro, and J. N. Onuchic, eLife **9**, e60312 (2020).
[5] J.-H. Su, P. Zheng, S. S. Kinrot, B. Bintu, and X. Zhuang, Cell **182**, 1641 (2020).
[6] A. B. Oliveira Junior, V. G. Contessoto, M. F. Mello, and J. N. Onuchic, Journal of Molecular Biology Diving into Chromatin across Space and Time, **433**, 166700 (2021).
[7] E. Dodero-Rojas, M. F. Mello, S. Brahmachari, A. B. Oliveira Junior, V. G. Contessoto, and J. N. Onuchic, Journal of Molecular Biology **435**, 10.1016/j.jmb.2023.168180 (2023), publisher: Academic Press.
[8] M. D. Pierro, D. A. Potoyan, P. G. Wolynes, and J. N. Onuchic, Proceedings of the National Academy of Sciences **115**, 7753 (2018).
[9] M. F. Mello, A. B. Oliveira Jr., E. Dodero-Rojas, S. Brahmachari, V. G. Contessoto, and J. N. Onuchic (2026), submitted.
[10] C. C. Correll, J. Bartek, and M. Dundr, Cells **8**, 869 (2019).
[11] H. D. Ou, S. Phan, T. J. Deerinck, A. Thor, M. H. Ellisman, and C. C. O'Shea, Science **357**, 10.1126/science.aag0025 (2017).
[12] J.-H. Su, P. Zheng, S. Kinrot, B. Bintu, and X. Zhuang, 10.5281/zenodo.3928890 (2020).
[13] Y. Luo, B. C. Hitz, I. Gabdank, J. A. Hilton, M. S. Kagda, B. Lam, Z. Myers, P. Sud, J. Jou, K. Lin, U. K. Baymuradov, K. Graham, C. Litton, S. R. Miyasato, J. S. Strattan, O. Jolanki, J.-W. Lee, F. Y. Tanaka, P. Adenekan, E. O'Neill, and J. M. Cherry, Nucleic Acids Research **48**, D882 (2020).
[14] W. Humphrey, A. Dalke, and K. Schulten, Journal of Molecular Graphics **14**, 33 (1996).
[15] John Stone, *An Efficient Library for Parallel Ray Tracing and Animation*, Ph.D. thesis, University of Missouri-Rolla (1998).
[16] V. Ramasubramani, B. D. Dice, E. S. Harper, M. P. Spellings, J. A. Anderson, and S. C. Glotzer, Computer Physics Communications **254**, 107275 (2020).
